# Mechanism of Tethered Agonist Binding to an Adhesion G-Protein-Coupled Receptor

**DOI:** 10.1101/2025.08.04.668378

**Authors:** Keya Joshi, Yinglong Miao

## Abstract

Adhesion GPCRs (ADGRs) contain a GPCR autoproteolysis-inducing (GAIN) domain that is proximal to the receptor N-terminus and undergoes autoproteolysis at a highly conserved GPCR proteolysis (GPS) site to generate the N-terminal fragment (NTF) and transmembrane C-terminal fragment (CTF). Dissociation of NTF reveals a peptide tethered agonist (TA) which is responsible for the activation of the ADGRs. The NTF-bound ADGRs contain the encrypted TA that assumes a β-strand configuration within the GAIN domain, which is markedly different from a U-shaped α-helical configuration of TA in the active cryo-EM structures of ADGRs. However, how the TA dramatically changes its configuration and binds to the ADGR CTF remains unknown. In this study, we have performed all-atom enhanced sampling simulations using a novel Peptide-Gaussian accelerated Molecular Dynamics (Pep-GaMD) method on TA binding to the ADGRD1. The Pep-GaMD simulations captured spontaneous binding of the TA into orthosteric pocket of the ADGRD1 and its large conformational transition from the extended β strand to the U-shaped α helical configuration. We were able to identify important low-energy conformations of the TA in the binding pathway, as well as different active and inactive states of ADGRD1, in the presence and absence of the Gs protein. Therefore, our Pep-GaMD simulations have revealed dynamic mechanism of the TA binding to an ADGR, which will facilitate rational design of peptide regulators of ADGRs.

**TOC Graphic:** 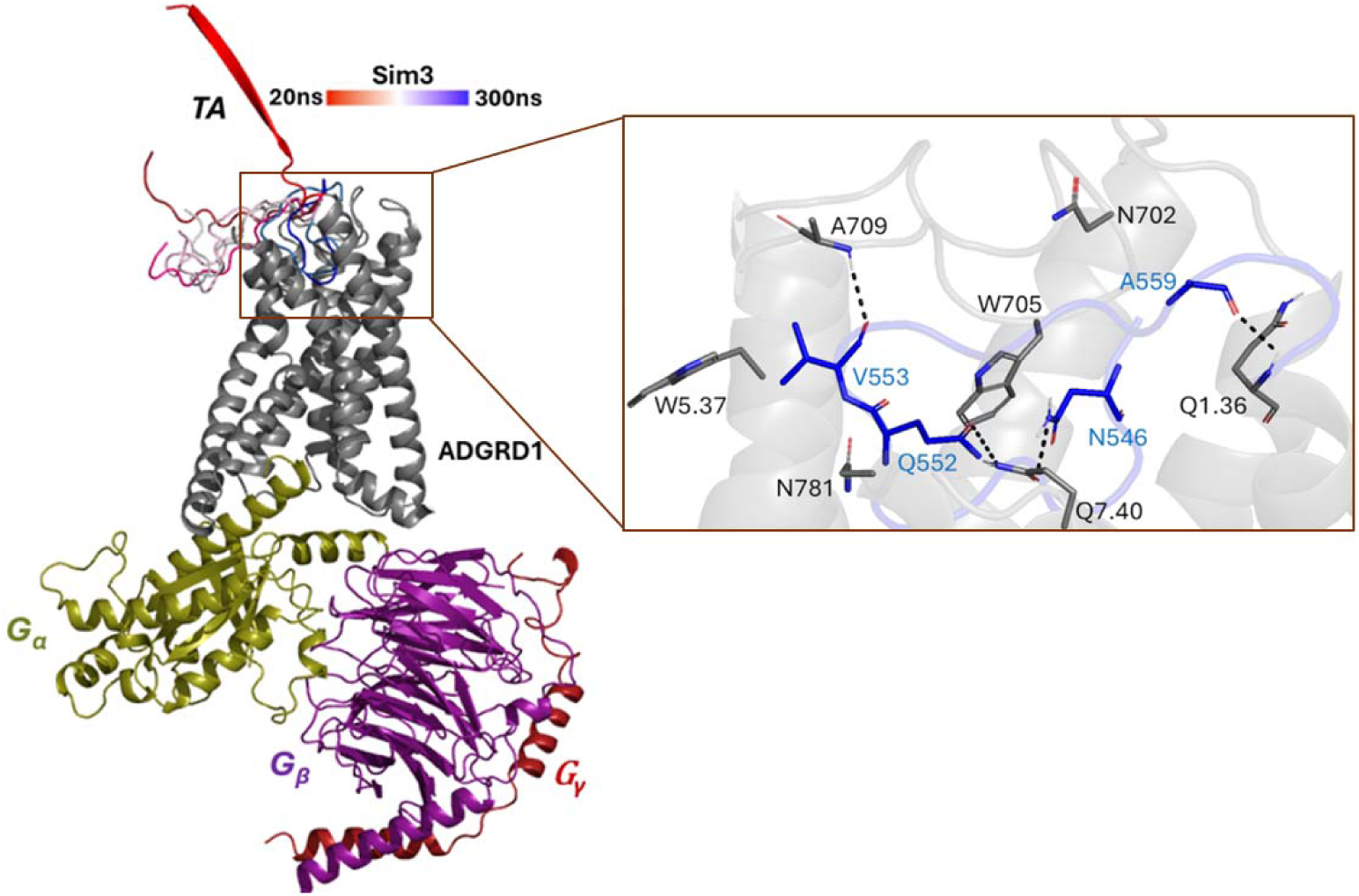

Peptide-Gaussian accelerated Molecular Dynamics (Pep-GaMD) simulations remarkably captured spontaneous binding of tethered agonist (TA) into the orthosteric pocket of adhesion G-protein–coupled receptor (ADGRD1) and large conformational transition of the TA from an extended β-strand to a U-shaped α-helical configuration. We were able to identify important low-energy conformations of the TA in the binding pathway and different states of the receptor from the Pep-GaMD simulations.

## 1. INTRODUCTION

G-protein-coupled receptors (GPCRs) are the largest family of human membrane proteins and play critical roles in cellular signaling and various physiological functions such as neurotransmission, sensory perception and hormonal regulation^1–4^. Therefore, GPCRs serve as primary targets for approximately 34% of FDA-approved marketed drugs^5–8^. Adhesion GPCRs (ADGRs) belong to the class B2 of GPCRs, comprising a total of 33 different members in humans^9–11^. ADGRs have several distinctive structural features that set them apart from other GPCRs - the presence of a large N-terminal extracellular domain (ECD) that comprises of several different cell-adhesion motifs including hormone binding domains, leucine rich repeats, immunoglobin and lectin binding domains. The ECD is involved in cell to cell or cell to matrix interactions^9, 12, 13^.

ADGRs play important roles in immune responses, tissue development and regulation of the cardiovascular and endocrine systems^14, 15^. ADGRD1 or GPR133 belongs to Group V class of ADGRs and is recognized as an oncogene in various cancers^16^. ADGRD1 and its immune-related genes are also recognized as potential biomarkers in non-small cell lung cancer (NSCLC)^17^. ADGRD1 activates G_S_-mediated signaling pathway for cytosolic cAMP. It has also been shown to be critical for the growth of glioblastoma, a brain malignancy^18^.

In 2022, the cryo-EM structure was reported for the ADGRD1 in complex with the G_S_ protein (PDB:7WU2)^16^. The first 7 amino acids located in the N-terminal region of the stalk with sequence “TNFAILM” acts as a tethered agonist (TA) of ADGRD1. The TA forms an alpha-helical conformation which bends nearly 180° downward into the orthosteric pocket and mediates important interactions with the transmembrane (TM) domain^16^. One such polar interaction is formed between the TA residue I549 and the extracellular loop 2 (ECL2) residue W705^ECL2^ which stabilizes the alpha-helical conformation of the TA^16^. The TA initiates the signal transduction through a direct contact with the ‘toggle switch’ residue W773^6.53^.

The TA which assumes a β-strand conformation within the hydrophobic core of the GPCR autoproteolysis-inducing (GAIN) domain undergoes a notable conformational rearrangement upon its dissociation from the GAIN^19^. Previous studies have reported the activation of ADGRs in the presence of tethered agonists^9, 16, 20–22^. However, the precise molecular mechanisms underlying binding of the TA in target receptor and the conformation rearrangement of the TA^20, 23^ remains widely unknown.

Building on Gaussian-accelerated Molecular Dynamics (GaMD)^24–28^, a new Pep-GaMD approach^29^ has been developed to explore the peptide-protein interactions, in which the essential potential energy of the peptide is selectively boosted to account for the peptide’s high flexibility and enhance peptide dissociation. In addition, another boost-potential is applied to the remaining potential energy of the entire system in a dual boost algorithm to enhance the peptide rebinding process. Dual-boost Pep-GaMD has captured repetitive peptide dissociation and binding events^30^ within significantly shorter simulation time (microsecond) compared to conventional molecular dynamics.

Here, we have applied Pep-GaMD simulations^29^ to simulate binding of the TA from an extended beta-strand conformation in the ADGRD1 receptor. The simulation findings have deepened our understanding of the TA binding and activation of ADGRs, which will facilitate rational design of peptide regulators of ADGRD1 and other ADGRs.

## 2. METHODS

### 2.1 System Setup

The cryo-EM structure of the active ADGRD1-G_S_ (PDB:7WU2) complex^16^ was used to set up the TA-bound ADGRD1-G_S_ simulation system **(Figure S1B)**. The first 7 amino acids with sequence “TNFAILM” act as a TA for ADGRD1, followed by linker residues “QVVPLEL” which covalently connects the TA to the receptor TM1 helix. Moreover, the G_S_ protein was removed to obtain only the TA-bound ADGRD1 simulation system **(Figure S1D)**.

Starting from the cryo-EM structure of TA-bound ADGRD1-Gs (PDB:7WU2) complex^16^, coordinates of the bound TA were replaced by the those of the AlphaFold^31^ model of the TA in the extended beta-strand conformation to obtain the simulation structure of TA-unbound ADGRD1 complex in the presence **(Figure S1A)** and absence **(Figure S1C)** of the G_S_-protein.

The four ADGRD1 simulation structures with TA bound **(Figure S1B and S1D)** and unbound conformations **(Figure S1A and S1C)** were embedded in a 1-palmitoyl-2-oleoyl-sn-glycero-3-phosphocholine (POPC) lipid bilayer using the CHARMM-GUI online server^32^. Residues at the protein termini were assigned neutral patches (acetyl and methyl amide). The N-termini of the TA (T545) was kept charged (NH3+). The systems were solvated in 0.15 M NaCl explicit solvent and immersed in a cubic TIP3P waterbox^33^ which was extended for 25 Å from the peptide−protein complex surface **(Figure S2A-B)**. The CHARMM36m force-field parameters^34^ were used for the proteins and lipids. The CHARMM-GUI^35^ output files and scripts were used for running MD simulations.

TA-bound ADGRD1-G_S_ system measured about 98.09 × 98.09 × 176.27 Å^3^ with 246 POPC lipid molecules (122 molecules on the upper leaflet and 124 molecules on the lower leaflet) and 37,449 water molecules, while TA-unbound ADGRG1-G_S_ system measured 114.21 × 114.21 × 202.34 Å^3^ with 348 POPC lipid molecules (172 molecules on the upper leaflet and 176 molecules on the lower leaflet) and 62,852 water molecules. Without the G_S_ protein, the TA-bound ADGRD1 system measured about 99.06 × 99.06 × 118.92 Å^3^ with 253 POPC lipid molecules (126 molecules on the upper leaflet and 127 molecules on the lower leaflet) and 23,543 water molecules, while the TA-unbound ADGRD1 system measured about 105.30 × 105.30 × 145.80 Å^3^ with 292 POPC lipid molecules (144 molecules on the upper leaflet and 148 molecules on the lower leaflet) and 35,982 water molecules.

### 2.2 Simulation Protocol

A total of 5000 steps of energy minimizations were carried out on the ADGRD1 systems, and a constant number, volume, and temperature (NVT) ensemble equilibration was performed for 125 ps at 310 K. Using an NPT ensemble, additional equilibration was carried out for 375 ps at 310 K. We then performed conventional MD (cMD) simulations on the four systems for 40 ns at 1 atm pressure and 310 K temperature with the AMBER22 software package^36, 37^. Long-range electrostatic interactions were computed with the particle mesh Ewald summation method, and a cutoff distance of 9 Å was used for the short-range electrostatic and Vander Waals interactions. After the 40 ns cMD runs, we performed Pep-GaMD^29^ equilibration for 50 ns for the four ADGRD1 (TA-bound ADGRD1-G_S,_ TA-unbound ADGRD1-G_S_, TA-bound ADGRD1 and TA-unbound ADGRD1) systems.

In order to observe the TA binding during the Pep-GaMD^29^ production simulations while keeping the boost potential as low as possible for accurate energetic reweighting, the (σ_0P_, σ_0D_) parameters were set to (3.0, 6.0 kcal/mol) with iEP set to 2 and iED set to 1 for the TA-unbound ADGRD1-G_S_ and TA-unbound ADGRD1 systems. This was followed by five independent Pep-GaMD^29^ production runs for 2500 ns for the TA-unbound ADGRD1-G_S_ and for 1000 ns for the TA-unbound ADGRD1 systems. Pep-GaMD^29^ production simulation frames were saved every 0.2 ps for analysis. The Pep-GaMD^29^ simulations on the TA-unbound ADGRD1 and TA-unbound ADGRD1-G_S_ systems are summarized in **Supplementary Tables 1 and 2**.

For the other two TA-bound ADGRD1-G_S_ and ADGRD1 systems, Pep-GaMD^29^ equilibration was performed with the default parameters (σ_0P_=6, σ_0D_=6, iEP=1, iED=1). This was followed by three independent Pep-GaMD^29^ production runs for 500 ns for each system. Pep-GaMD^29^ production simulation frames were saved every 0.2 ps for analysis. The Pep-GaMD^29^ simulations on the TA-bound ADGRD1 and TA-bound ADGRD1-G_S_ systems are summarized in **Supplementary Tables 3 and 4**.

### 2.3 Simulation Analysis

Simulation analysis was carried out using CPPTRAJ^38^ and VMD^39^. The software tools were applied to track binding of the TA in the ADGRD1 receptor. Fraction of the native contacts between the TA and receptor and the distance between the NE1 atom of W714^5.37^ and O atom of L708^ECL2^ were selected to calculate the 2D free energy profiles to characterize the TA binding pathway.

In addition, fraction of the native contacts between the TA and receptor and the distance between the Cα atoms of V661^3.58^ and K760^6.40^ were also selected to calculate the 2D free energy profiles to characterize the TA-induced conformational transitions in the ADGRD1 receptor. 2D free energy profiles were reweighted using the PyReweighting toolkit^26^. A bin size of 0.1 % was used for the fraction of native contacts between the TA and receptor while a bin size of 1 Å was used for the W714^5.37^-L708^ECL2^ and V661^3.58^-K760^6.40^ distances. The cutoff was set to 500 frames in one bin for reweighting. The Density Peak (DPeak) clustering algorithm^40^ was used to cluster snapshots of the receptor and TA conformations with all Pep-GaMD^29^ production simulations combined for each system. The Pep-GaMD^29^ simulation snapshots of the TA in the ADGRD1 were clustered to identify the representative low-energy conformations shown in the 2D free energy profiles.

## 3. RESULTS AND DISCUSSION

### 3.1 Pep-GaMD simulations captured spontaneous binding of the TA into orthosteric site of the ADGRD1 receptor

In four out of five independent 2500 ns Pep-GaMD^29^ production simulations (Sim1 to Sim4), complete binding of the TA was observed from the extracellular region to the orthosteric pocket in the ADGRD1 receptor at ∼500-800 ns, for which fraction of the native contacts between the TA and receptor relative to the cryo-EM TA-bound conformation^16^ increased to 0.9-1.0 **(Figure 1A and 1B)**. During binding, the TA moved from the extracellular solvent into the orthosteric pocket of the receptor through the receptor’s extracellular mouth between ECL2, ECL3, TM5, TM6 and TM7. The TA also underwent dramatic conformational transition from an extended beta-strand to a U-shaped alpha-helical conformation at the orthosteric site **(Figure 1A)**. In Sim5, the TA showed only partial binding to the ADGRD1 receptor with interactions with the ECL3, for which fraction of the native contacts between the TA and receptor relative to the cryo-EM conformation reached only ∼0.3-0.5 **(Figure 1B)**.

**Figure 1.**
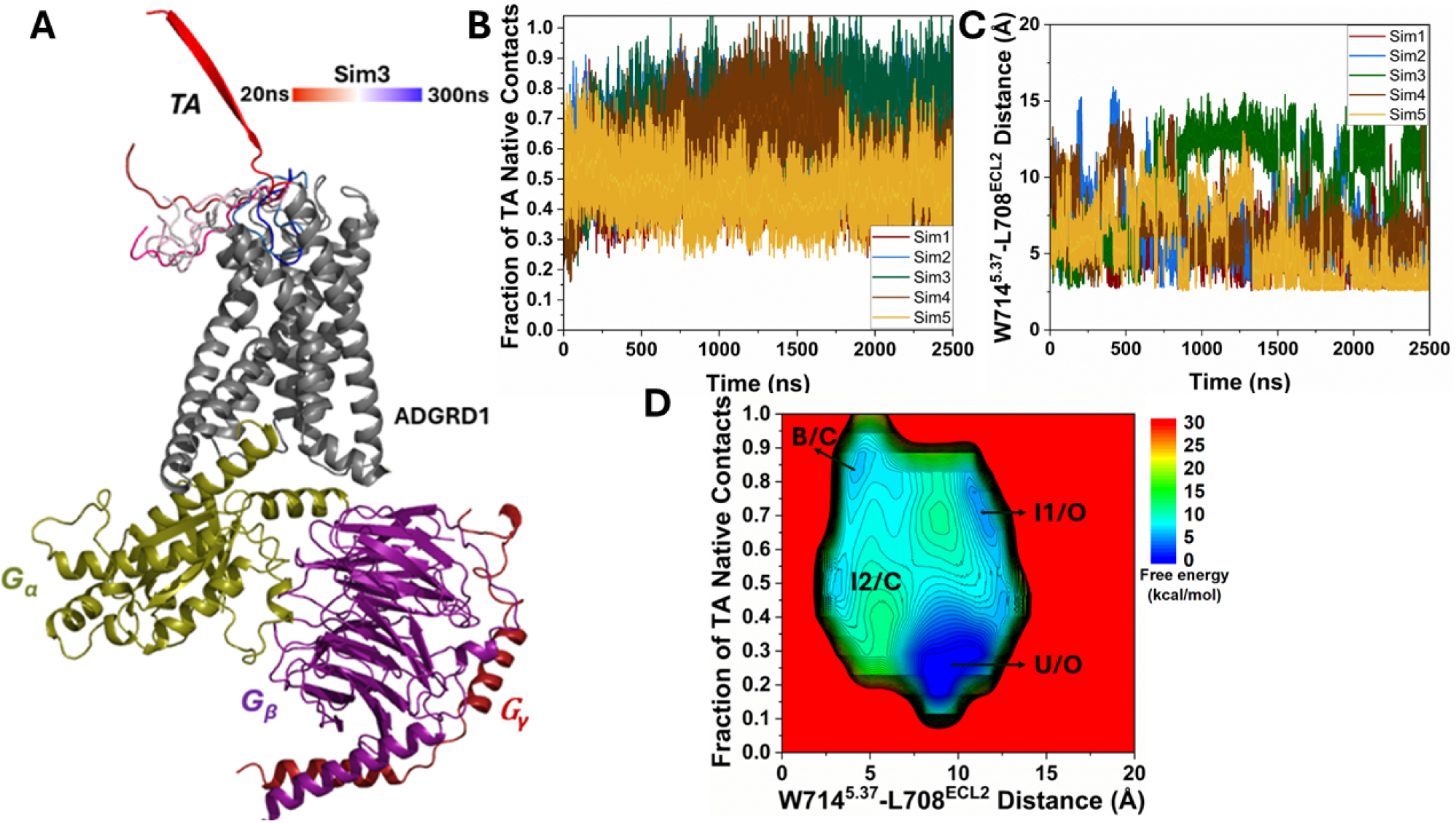
Binding of tethered agonist (TA) in the G_S_ protein-coupled ADGRD1 receptor was observed in Pep-GaMD simulations: **(A)** A representative binding pathway of the TA, which is colored by simulation time. **(B)** Time course of the fraction of native contacts between the TA and receptor calculated from five 2500 ns Pep-GaMD simulations. **(C)** Time course of the distance between the NE1 atom of W714^5.37^ and O atom of L708^ECL2^ calculated from five 2500 ns Pep-GaMD simulations. **(D)** 2D potential of mean force (PMF) free energy profile regarding the fraction of native contacts between the TA and receptor and W714^5.37^-L708^ECL2^ distance calculated by combining five 2500 ns Pep-GaMD simulations. The low-energy states are labeled as “Unbound/Open” (U/O), “Intermediate 1/Open” (I1/O), “Intermediate 2/Closed” (I2/C) and “Bound/Closed” (B/C).

The five Pep-GaMD^29^ simulations showed similar boost potentials with averages of ∼37.43-40.64 kcal/mol and standard deviations (SDs) of ∼5.35-5.79 kcal/mol **(Table S2)**. We combined all the Pep-GaMD^29^ production simulations to calculate free energy profiles for binding of the TA in the ADGRD1 through energetic reweighting^26^ **(See Methods).** In the 2D free energy profile of the fraction of native contacts between the TA and receptor and the distance between residues W714^5.37^-L708^ECL2^ **(Figure 1C)**, we identified four low-energy conformational states, i.e., the “Unbound/Open” (U/O), “Intermediate 1/Open” (I1/O), “Intermediate 2/Closed” (I2/C) and “Bound/Closed” (B/C) **(Figure 1D)**.

In the “Unbound/Open” (U/O) state, the TA adopted an extended beta-strand conformation pointing away from the orthosteric site **(Figure 1A)**. This conformation represented the unbound state when the TA was embedded within the β-sheet core of the GAIN domain^21^. In the U/O state, the fraction of native contacts between the TA and receptor was only ∼0.1-0.2 **(Figure 1B)** while the distance between the NE1 and O atoms of residues W714^5.37^-L708^ECL2^ was 10 Å **(Figure 1C)**, which indicates the open or expanded state of the ADGRD1 pocket when no TA is bound to the orthosteric site **(Figure 1D)**.

In the “Intermediate 1/Open” (I1/O) state, the TA binds to the extracellular mouth of the receptor with the fraction of native contacts between the TA and receptor maintained around ∼0.5-0.7 **(Figure 1B)** while the distance between the NE1 and O atoms of residues W714^5.37^-L708^ECL2^ remains centered around 10 Å **(Figure 1C)**, which again indicates that the receptor pocket was in an open or expanded state when no TA was bound to the orthosteric site **(Figure 1D)**.

In the “Intermediate 2/Closed” (I2/C) state, the TA enters the orthosteric pocket with the fraction of native contacts between the TA and receptor centered around ∼0.5-0.6 **(Figure 1B)** while the distance between the NE1 and O atoms of residues W714^5.37^-L708^ECL2^ decreased to ∼2.5-3 Å from 10 Å **(Figure 1C)**, which indicates that as the TA binds into the orthosteric site, the interaction with the pocket residues increases and the pocket space gets more compacted. Finally, the TA samples the “Bound/Closed” (B/C) state, in which the fraction of native contacts between the TA and receptor was ∼0.8-0.9 **(Figure 1B)**, being similar to the cryo-EM structure while the distance between the NE1 and O atoms of residues W714^5.37^-L708^ECL2^ centered around ∼2.5-3 Å **(Figure 1C)**.

### 3.2 TA exhibited flexibility in the ADGRD1-G_S_ protein complex

In three independent 500 ns Pep-GaMD production simulations of TA-bound ADGRD1-G_S_ protein complex, the bound TA showed flexibility within the orthosteric pocket in the ADGRD1 receptor. The system sampled a new “Intermediate 2/Closed” (I2/C) conformation alongside the “Bound/Closed” (B/C) conformation **(Figure 2A)**. The three Pep-GaMD^29^ simulations on the TA-bound ADGRD1-G_S_ protein complex showed boost potentials with averages of ∼12.38-13.28 kcal/mol and SDs of ∼3.87-4.04 kcal/mol **(Table S4)**.

**Figure 2.**
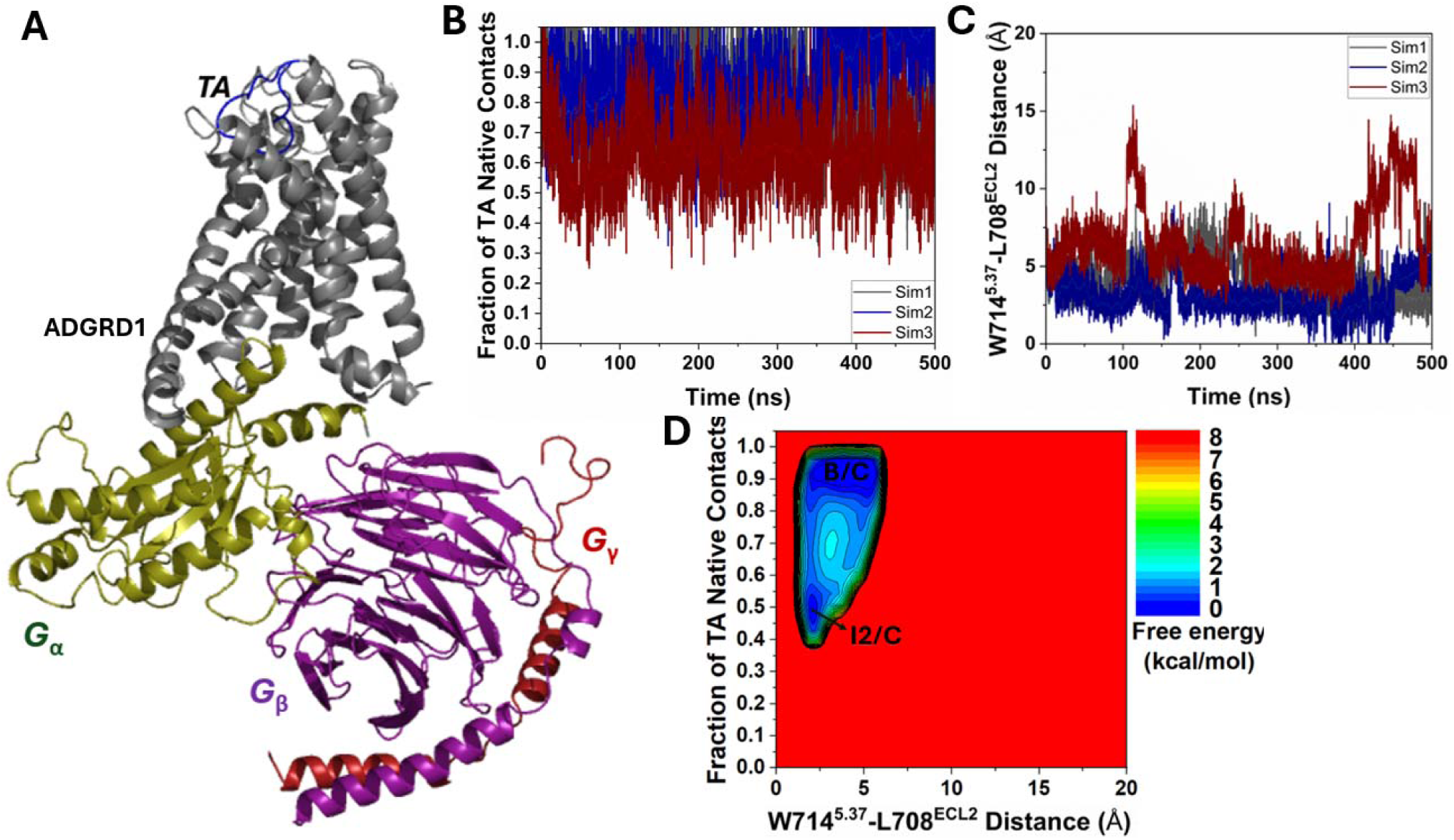
Dynamic motions of bound tethered agonist (TA) in the ADGRD1-G_S_ protein complex were observed in Pep-GaMD simulations: **(A)** A representative structure of ADGRD1-G_S_ protein complex depicting the flexibility of bound TA. **(B)** Time course of the fraction of native contacts between the TA and receptor calculated from three 500 ns Pep-GaMD simulations. **(C)** Time course of the distance between the NE1 atom of W714^5.37^ and O atom of L708^ECL2^ calculated from three 500 ns Pep-GaMD simulations. **(D)** 2D potential of mean force (PMF) free energy profile regarding the fraction of native contacts between the TA and receptor and W714^5.37^-L708^ECL2^ distance calculated by combining three 500 ns Pep-GaMD simulations. The low-energy states are labeled as “Intermediate 2/Closed” (I2/C) and “Bound/Closed” (B/C).

We combined all three Pep-GaMD^29^ production simulations on the TA-bound ADGRD1-G_S_ complex to calculate free energy profiles through energetic reweighting^26^ **(See Methods).** In the 2D free energy profile of the fraction of native contacts between the TA and receptor and the W714^5.37^-L708^ECL2^ distance **(Figure 2D)**, we identified two low-energy conformational states, i.e., the “Intermediate 2/Closed” (I2/C) and “Bound/Closed” (B/C). In the “I2/C” state, the fraction of native contacts between the TA and receptor pocket was ∼0.5 **(Figure 2B)**, while the distance between W714^5.37^-L708^ECL2^ centered around ∼2.5 Å **(Figure 2C)**.

Both the “I2/C” and “B/C” states were also observed in the Pep-GaMD^29^ simulations of TA-unbound ADGRD1-G_S_ system (**Figure 1D**), indicating the presence of an intermediate conformation of the TA in the receptor orthosteric pocket (“I2”) before it reached the final bound conformation as in the cryo-EM structure. This intermediate conformation also accounts for the TA’s dynamic motions within the receptor orthosteric pocket.

### 3.3 Low-energy conformational states of TA binding in the ADGRD1 receptor

The DPeak clustering algorithm^40^ was used to cluster snapshots of the ADGRD1 receptor with all the Pep-GaMD^29^ production simulations combined to identify the representative low-energy conformations of the TA in the TA-unbound ADGRD1-G_S_ simulation system **(Figure 3)**.

**Figure 3.**
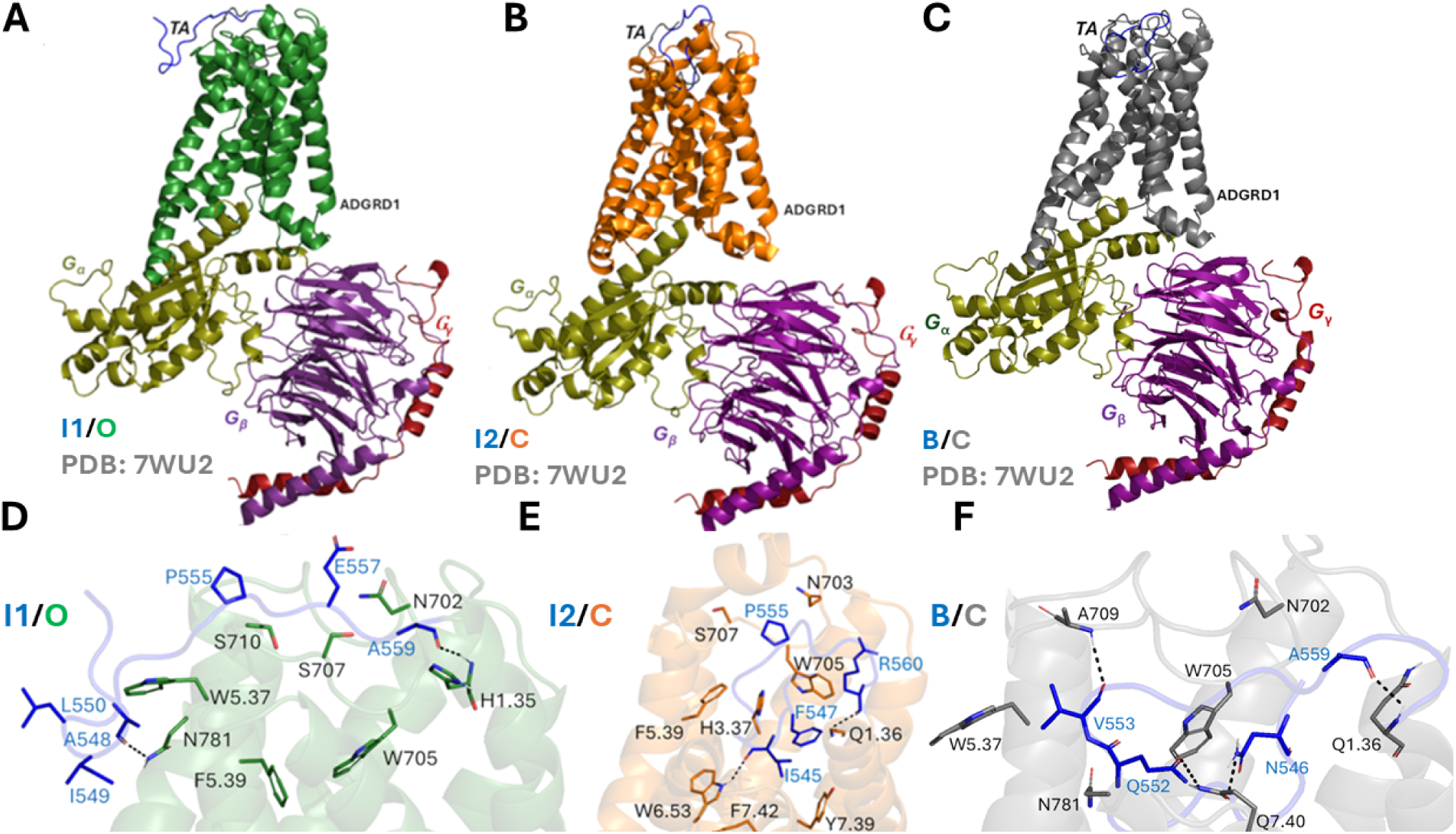
Low-energy conformational states of the tethered agonist (TA) binding in the ARGRD1-G_S_ complex: **(A-C)** The “Intermediate 1/Open” (I1/O), “Intermediate 2/Closed” (I2/C) and “Bound/Closed” (B/C) states. The cryo-EM bound conformation of TA (PDB: 7WU2) is shown in grey as reference. **(D)** Critical residue interactions between TA (blue) and ADGRD1 (green) observed in the “I1/O” state. The TA formed polar and hydrophobic interactions with receptor residues H^1.35^, N702^ECL2^, W705^ECL2^, S707^ECL2^, S710^ECL2^, W^5.37^, F^5.39^ and N781^ECL3^. **(E)** Critical residue interactions between TA (blue) and ADGRD1 (orange) observed in the “I2/C” state. The TA formed polar and hydrophobic interactions with receptor residues Q^1.36^, H^3.37^, N703^ECL2^, W705^ECL2^, S707^ECL2^, F^5.39^, W^6.53^, Y^7.39^ and F^7.42^. **(F)** Critical residue interactions between TA (blue) and ADGRD1 (grey) observed in the “B/C” state. The TA formed polar and hydrophobic interactions with receptor residues Q^1.36^, N702^ECL2^, W705^ECL2^, A709ECL2, W5.37, N781ECL3 and Q7.40.

In the “Intermediate 1/Open” (I1/O) state, the TA which was initially pointing away in the U/O state binds near the extracellular mouth (ECL3, TM5, TM6 and TM7) of the ADGRD1 receptor **(Figure 3A)**. The tethered agonist residues (A548 and A559) formed two polar interactions with receptor residues H^1.35^ and N781^ECL3^ in the I1/O state. Strong hydrophobic interactions were also observed between the TA and the receptor residues N702^ECL2^, W705^ECL2^, S707^ECL2^, S710^ECL2^, W^5.37^ and F^5.39^ **(Figure 3A and 3D)**.

In the “Intermediate 2/Closed” (I2/C) state, the TA enters the orthosteric pocket through the extracellular mouth (ECL2, ECL3, TM5, TM6 and TM7) of the receptor. The orientation of the TA in the I2/C state is different compared to its bound state-The linker (“QVVPLEL”) connecting the TA and TM1 shows higher flexibility and also the N-terminal part of the TA points upwards compared to its bound conformation **(Figure 3B)**. In the I2/C state, the TA residues F547 and P555 formed strong hydrophobic interactions with the pocket residues H^3.37^, W705^ECL2^, F^5.39^, Y^7.39^ and F^7.42^ while residues R560 and I545 formed polar interactions with receptor residues Q^1.36^ and W^6.53^ **(Figure 3E)**. I545 in the TA was in direct contact with the putative “toggle” switch W773^6.53^, which is known to be the activation hub of many GPCRs^16^ **(Figure 3E)**. Alteration of the conformation of W773^6.53^ could directly modulate ADGRD1 activity. It has also been observed that the TA mediates the signal transduction by interacting with the “toggle” switch residue^16^. This highly conserved bulky residue tethers TM3, TM5 and TM6 helices by forming a hydrophobic core with F547, M551 and I549 residues from the TA.

As the TA binds to the orthosteric site in the “Bound/Closed” (B/C) state, it undergoes the conformational transition from an extended beta-strand to an alpha-helical conformation^21^, which is stabilized by receptor residues from TM1, TM2, TM5, TM6, TM7, ECL3 and ECL2 **(Figure 3C)**. The bound conformation of the TA adopts two rim states-Lower rim (“TNFAILM”) and Upper rim (“QVVPLEL”). The lower rim of the TA assumed an alpha-helical conformation, deeply inserted into the hydrophobic core of the receptor and showed interactions with receptor residues W705^ECL2^, W^5.37^, N781^ECL3^ and Q^7.40^ **(Figure 3F)**. Previous studies^16^ have experimentally validated W705^ECL2^ residue in ECL2 to be critical for the activation of ADGRD1 by the tethered agonist. Substantial reduction of the agonistic potency of the tethered agonist was observed when W705^ECL2^ was mutated to alanine^16^. The upper rim of the TA forms two polar interactions with receptor residue A709^ECL2^ and Q^1.36^ **(Figure 3F)**. Ala substitution of Q^1.36^ in ADGRD1 significantly impaired the basal activity of ADGRD1.

### 3.4 Pep-GaMD simulations captured partial binding of TA to the ADGRD1 receptor in the absence of the G_S_-protein

In five independent 1000 ns Pep-GaMD^29^ production simulations of the TA-unbound ADGRD1 system, partial binding of the TA was observed from the extracellular region to the ECL3 on the ADGRD1 receptor surface at ∼300-500 ns **(Figure 4A)**, for which the majority of the fraction of native contacts between the TA and receptor relative to the cryo-EM bound conformation reached ∼0.5-0.6 **(Figure 4B and 4D)**. The five Pep-GaMD^29^ simulations showed similar boost potentials with averages of ∼18.62-19.33 kcal/mol and SDs of ∼4.44-4.52 kcal/mol **(Table S1)** while the three Pep-GaMD^29^ simulations on the TA-bound ADGRD1 system showed boost potentials with averages of ∼12.24-12.76 kcal/mol and standard deviations (SDs) of ∼3.90-3.95 kcal/mol **(Table S3)**. We combined all the Pep-GaMD^29^ production simulations to calculate the free energy profiles through energetic reweighting^26^ **(See Methods)**.

**Figure 4.**
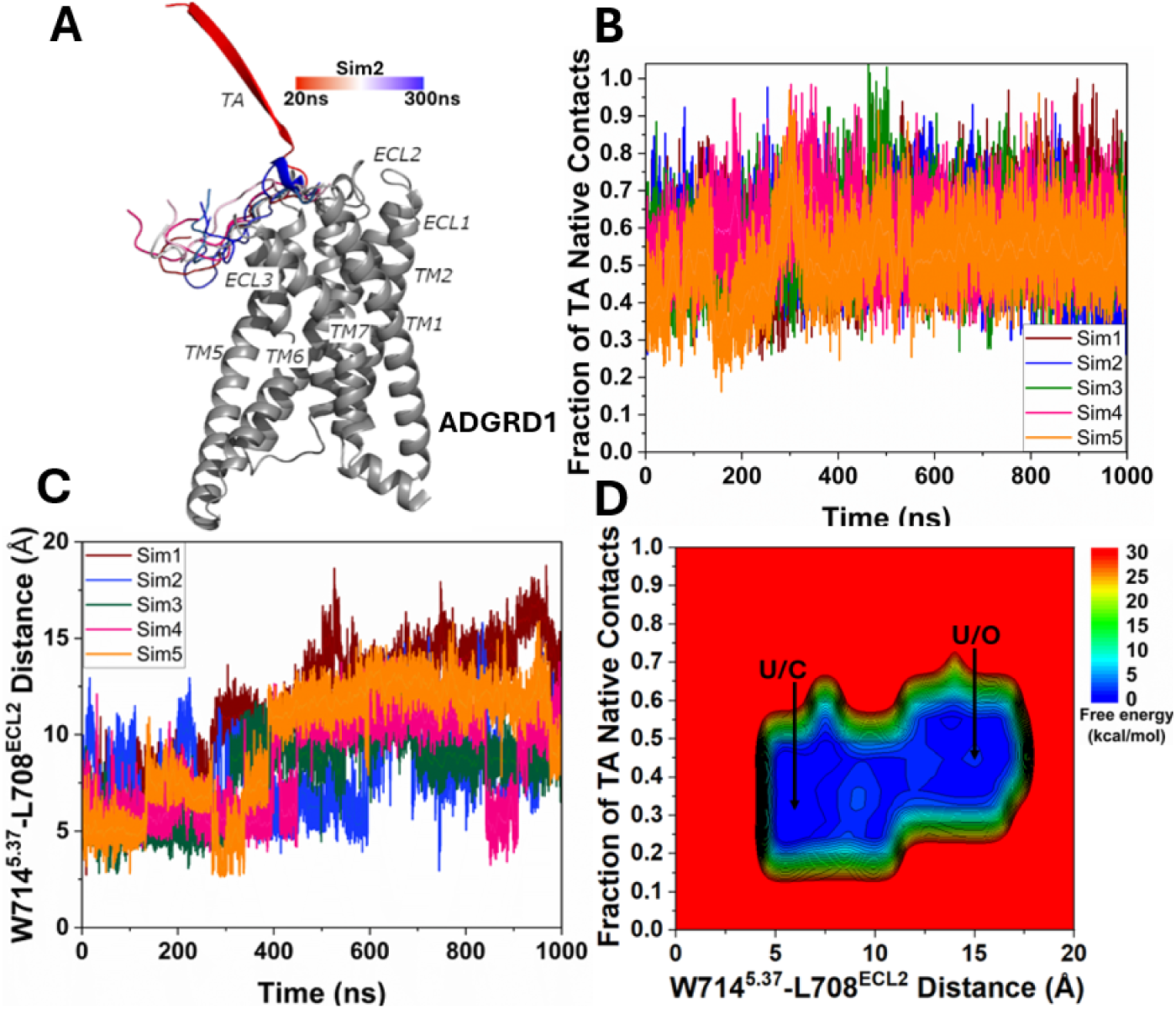
Partial binding of tethered agonist (TA) in the ADGRD1 receptor without the G_S_ protein was observed in Pep-GaMD simulations: **(A)** A representative partial binding pathway of the TA, which is colored by simulation time. **(B)** Time course of the fraction of native contacts between the TA and receptor calculated from five 1000 ns Pep-GaMD simulations. **(C)** Time course of the distance between the NE1 atom of W714^5.37^ and O atom of L708^ECL2^ calculated from five 1000 ns Pep-GaMD simulations. **(D)** 2D potential of mean force (PMF) free energy profile regarding the fraction of native contacts between the TA and receptor and W714^5.37^-L708^ECL2^ distance calculated by combining five 1000 ns Pep-GaMD simulations. The low-energy states are labeled as “Unbound/Closed” (U/C) and “Unbound/Open” (U/O).

In the 2D free energy profile of the fraction of native contacts between the TA and receptor and the distance between W714^5.37^-L708^ECL2^ **(Figure 4D)**, we identified two low-energy conformational states, i.e., the “Unbound/Close” (U/C) and “Unbound/Open” (U/O). In the “Unbound/Close” (U/C) state, the TA adopted an extended beta-strand conformation pointing away from the orthosteric site **(Figure 4A)**, where the fraction of native contacts between the TA and receptor was ∼0.2-0.4 **(Figure 4B and 4D)**, while the distance between residues W714^5.37^-L708^ECL2^ was ∼5 Å **(Figure 4C and 4D)**. In the “Unbound/Open” (U/O) state, the TA bound near the extracellular mouth around ECL3 on the ADGRD1 receptor surface **(Figure 4A)**, where the fraction of native contacts between the TA and receptor reached ∼0.3-0.4 **(Figure 4B and 4D)**, while the distance between residues W714^5.37^-L708^ECL2^ at the receptor extracellular mouth increased to ∼10-15 Å **(Figure 4C and 4D)**.

### 3.5 ADGRD1 transitioned to the inactive state upon removal of the G_S_-protein and partial binding of the TA

In three independent 500 ns Pep-GaMD^29^ simulations on TA-bound ADGRD1-G_S_ complex, the ADGRD1 receptor maintained the active state and sampled two low-energy conformational states, i.e., the “Intermediate 2/Active” (I2/A) and “Bound/Active” (B/A) **(Figure S4B)**. In both the I2/A and B/A states, the distance between the Cα atoms of V661^3.58^ and K760^6.40^ residues was centered around ∼15-16 Å **(Figure S4A)**. Similarly, in the Pep-GaMD simulations on TA-unbound ADGRD1-G_S_ complex, the ADGRD1 receptor maintained the active state throughout the simulations and sampled two low-energy conformational states, i.e., the “Unbound/Active” (U/A) and “Bound/Active” (B/A) **(Figure S3B)** with the distance between the Cα atoms of V661^3.58^ and K760^6.40^ residues centered around ∼15-16 Å **(Figure S3A)**. These findings conclude that in the presence of G_S_-protein and bound TA, the receptor maintains an active state.

Interestingly, ADGRD1 receptor was also able to maintain its active state during the Pep-GaMD^29^ simulations on the TA-bound ADGRD1 complex and sampled two low-energy conformational states, i.e., the “Intermediate 2/Active” (I2/A) and “Bound/Active” (B/A) **(Figure S5C)**. In both the I2/A and B/A states, the receptor maintained its active state with V661^3.58^-K760^6.40^ distance centered around ∼15-16 Å **(Figure S5B)**. In the I2/A state, the TA residues (T545, N546, I549, Q552 and V553) also formed polar interactions with the pocket residues N702^ECL2^, W705^ECL2^, A709^ECL^, W^5.37^ and V^6.57^ as highlighted in **Figure S6A and S6B**.

In contrast, deactivation of the ADGRD1 receptor was observed in the Pep-GaMD^29^ simulations of the TA-unbound ADGRD1 system in the absence of the G_S_ protein. In 2D free energy profile of the fraction of native contacts between the TA and receptor and the distance between V661^3.58^ and K760^6.40^ **(Figure 5A and 5B)**, we identified two low-energy conformational states, i.e., the “Unbound/Active” (U/A) and “Unbound/Inactive” (U/IN). In the “Unbound/Active” (U/A) state, the TA remained in the extended unbound conformation and the receptor maintained its active state with the V661^3.58^-K760^6.40^ distance centered around ∼16Å **(Figure 5B)**. In the “Unbound/Inactive” (U/IN) state, the TA moved close to the extracellular mouth near ECL3 on the receptor surface **(Figure 5C)** while the receptor became deactivated with the V661^3.58^-K760^6.40^ distance decreased to ∼10 Å compared to ∼16 Å distance in the active cryo-EM structure **(Figure 5D)**. These findings highlight that in the absence of G_S_-protein and partial binding of TA; the receptor becomes inactive.

**Figure 5.**
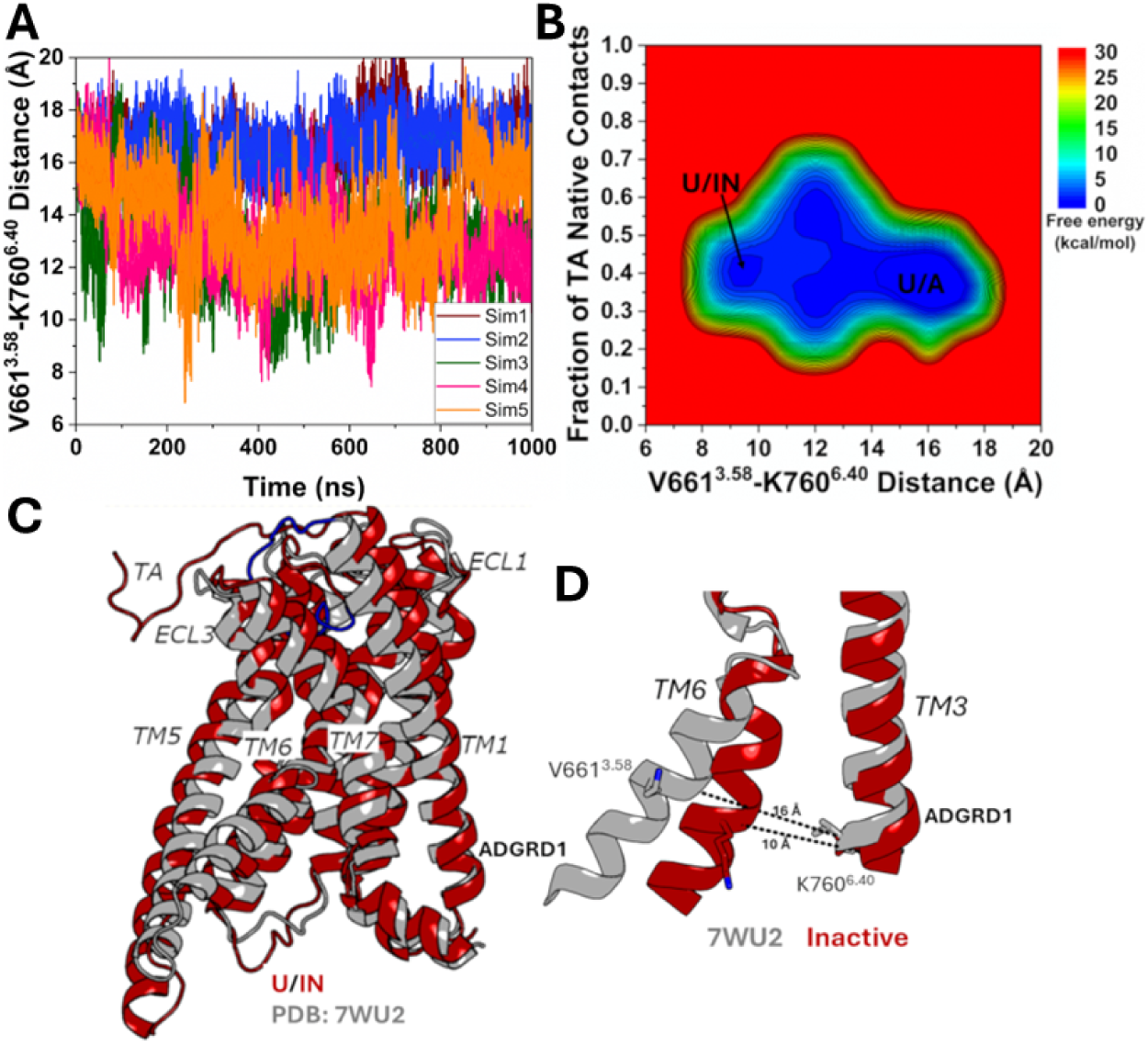
Inactivation of the ADGRD1 receptor was observed during only partial binding of tethered agonist (TA) without the G_S_ protein in Pep-GaMD simulations: **(A)** The distance between the Cα atoms of V661^3.58^ and K760^6.40^ calculated from five 1000 ns Pep-GaMD simulations. **(B)** 2D potential of mean force (PMF) free energy profile regarding the fraction of native contacts between the TA and receptor and V661^3.58^-K760^6.40^ distance calculated by combining five 1000 ns Pep-GaMD simulations. The low-energy states are labeled as “Unbound/Active” (U/A) and “Unbound/Inactive” (U/IN). **(C)** The “Unbound/Inactive” (U/IN) conformation. The cryo-EM bound conformation of TA (PDB: 7WU2) is shown in grey as reference. **(D)** Comparison of the active cryo-EM conformation (grey) with the Inactive conformational state of the ADGRD1 receptor observed in the Pep-GaMD simulations.

## 4. CONCLUSIONS

In this study, all-atom Pep-GaMD^29^ simulations have been applied to simulate binding of the TA from an extended beta-strand conformation outside of the ADGRD1 receptor to a U-shaped alpha-helical conformation in the receptor’s orthosteric pocket^16^. Pep-GaMD^29^ simulations captured distinct low-energy states of the TA along the binding pathway, as well as low-energy conformations of the ADGRD1 in the active and inactive states.

The TA assumed a β-strand configuration **(Figure 1A and 4A)** within the hydrophobic core of GAIN domain. It is distinct from the observed U-shaped configuration in the bound active cryo-EM structure^9^. Mutations disrupting the hydrophobic packing between the TA and the surrounding GAIN domain residues could lead to release of the TA from the GAIN domain and binding to the receptor TM domain, leading to the activation of the ADGRs^21^.

The Pep-GaMD^29^ simulations revealed two novel intermediate conformations of the TA - “Intermediate 1” (I1) and “Intermediate 2” (I2) - before the TA reached the final bound state as observed in the cryo-EM structure^16^. In the I1 state, the TA bound to the extracellular mouth near ECL3, TM5, TM6 and TM7 of the receptor **(Figure 3A)**. In the I2 state, the TA entered the orthosteric binding pocket, although the N terminus of the TA pointed away from the target site and the linker residues “QVVPLEL” connecting the TA and TM1 showed high flexibility **(Figure 3B)**.

The TA in the “Bound/Active” (B/A) state was stabilized by residues from the orthosteric pocket **(Figure 3C and 3F)**. Mutation of residues present in the TA or the interacting residues from the receptor’s orthosteric pocket significantly impaired the basal activity of ADGRD1. Previous studies^16^ have experimentally validated residue W705 in ECL2 to be critical for activation of the ADGRD1.

It is important to note that the presence of the G_S_ protein and binding of the TA are critical to maintain the ADGRD1 receptor in the active state. The receptor became inactive upon removal of the G_S_ protein and only partial binding of the TA in the Pep-GaMD simulations.

In summary, we have uncovered the pathway and dynamic mechanism of TA binding to an ADGR using all-atom Pep-GaMD^29^ simulations. The understanding we gained regarding the binding of the highly flexible TA and its dramatic conformational transition in the receptor provides valuable insights into rational design of novel peptide regulators of ADGRs.

## ASSOCIATED CONTENT

### Supporting Information

The Supporting Information includes details about the Pep-GaMD method, Tables S1 to S4 and Figures S1 to S6. This information is available free of charge via the Internet at http://pubs.acs.org.

We have also shared all the necessary simulation input files for Pep-GaMD simulations of the TA-unbound ADGRD1, TA-unbound ADGRD1-G_S_, TA-bound ADGRD1 and TA-bound ADGRD1-G_S_ systems in the public repository Figshare with the following links- https://doi.org/10.6084/m9.figshare.29691206

https://doi.org/10.6084/m9.figshare.29689628

https://doi.org/10.6084/m9.figshare.29682797

https://doi.org/10.6084/m9.figshare.29682371

## AUTHOR INFORMATION

### Author contributions

Y.M. designed the research; K.J. performed research; K.J. and Y.M. analyzed data; and K.J. and

Y.M. wrote the paper.

### Notes

The authors declare no competing financial interest.

## Supporting information

The Supporting Information includes details about the Pep-GaMD method, Tables S1 to S4 and Figures S1 to S6

## ACKNOWLEDGMENTS

This work used supercomputing resources with allocation award TG-MCB180049 through the Extreme Science and Engineering Discovery Environment ACCESS, which is supported by National Science Foundation grant number ACI-1548562, and project M2874 through the National Energy Research Scientific Computing Center (NERSC), which is a U.S. Department of Energy Office of Science User Facility operated under Contract No. DE-AC02-05CH11231. This work was also supported by the National Institutes of Health (R01GM132572) and the startup funding project 27110 at the University of North Carolina-Chapel Hill.

---

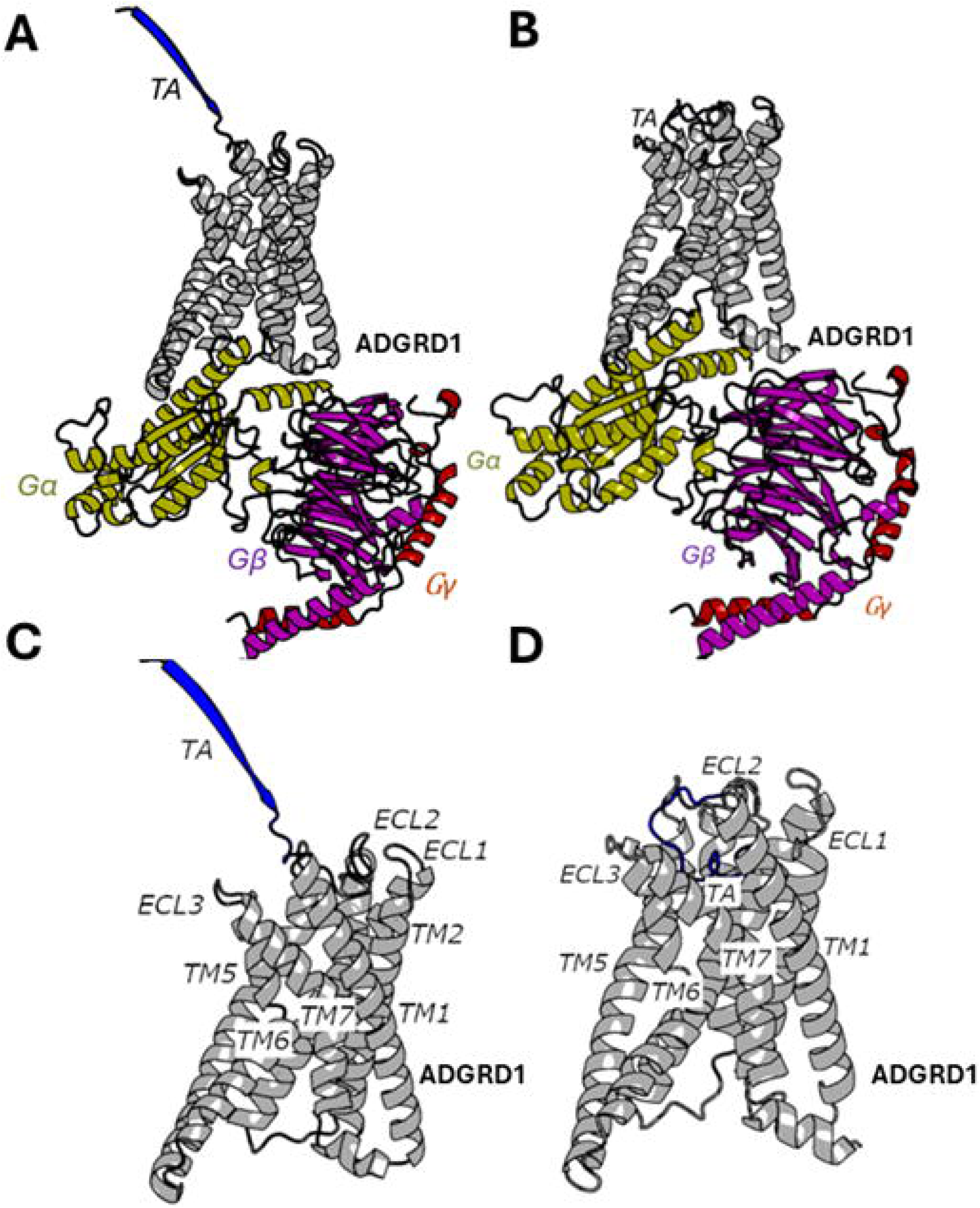

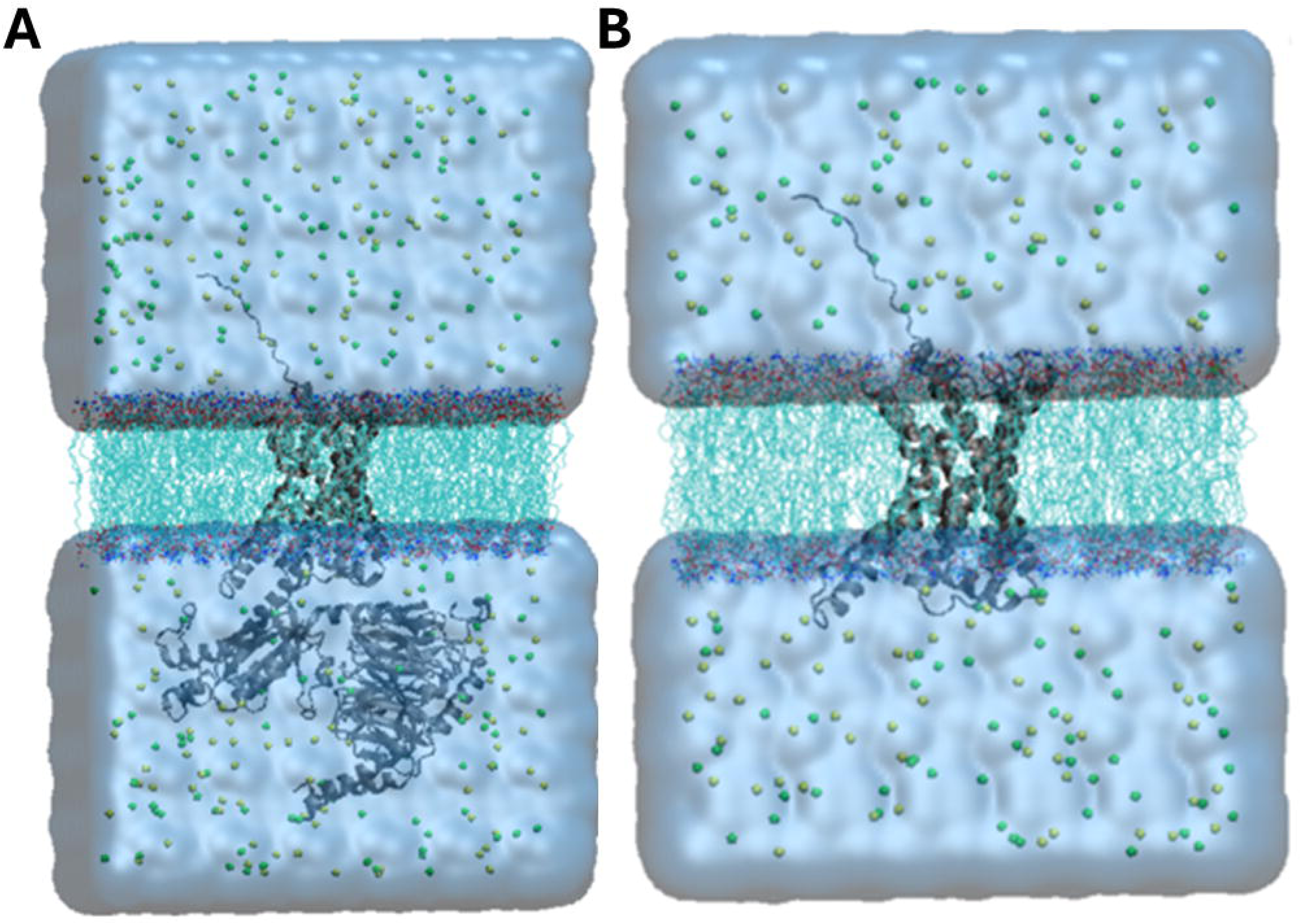

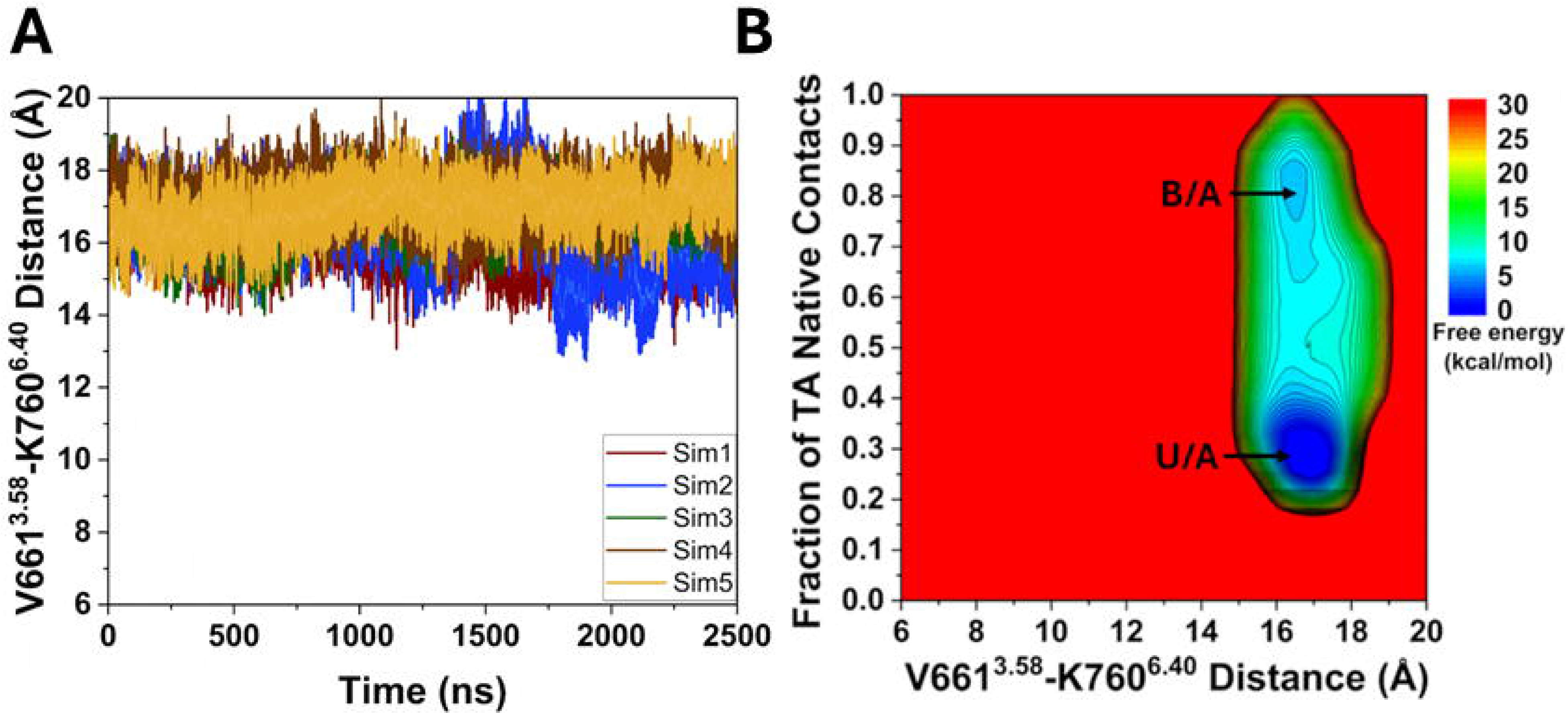

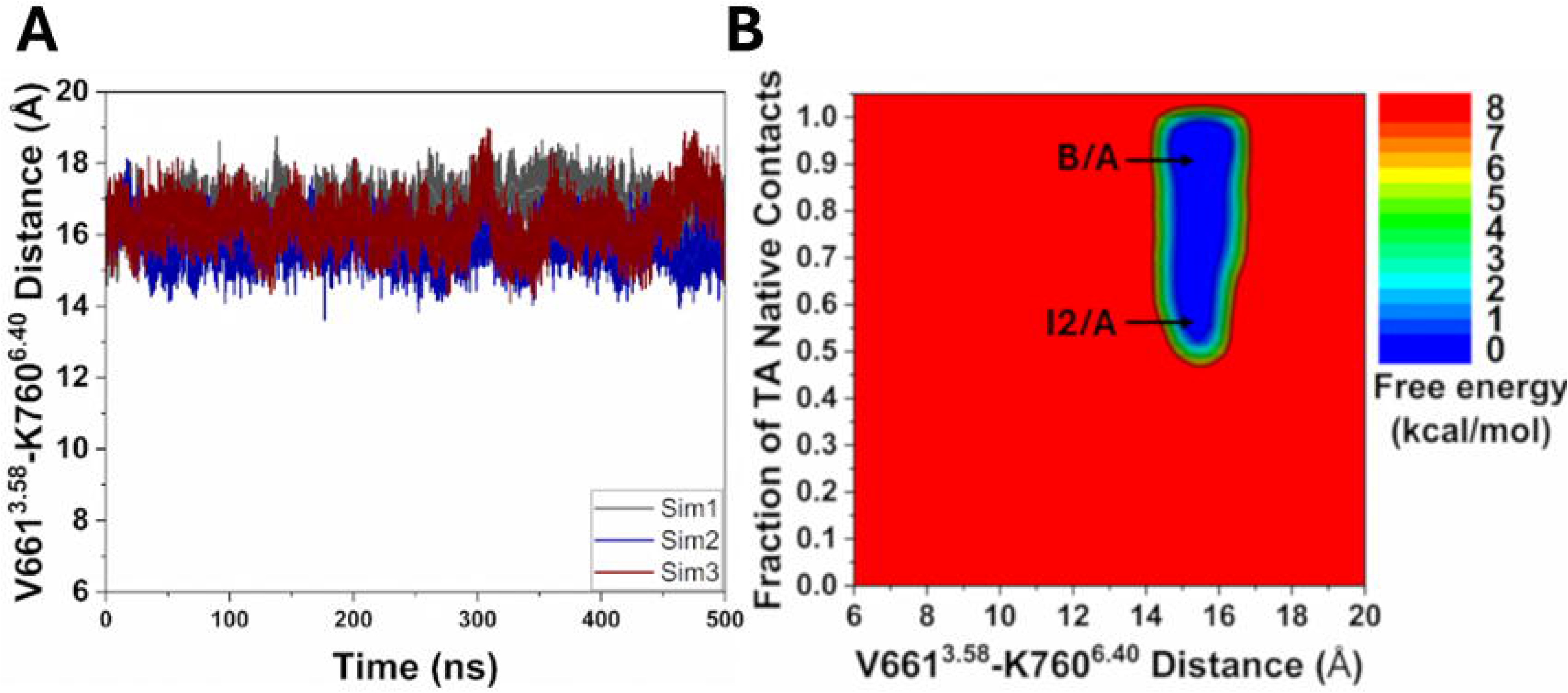

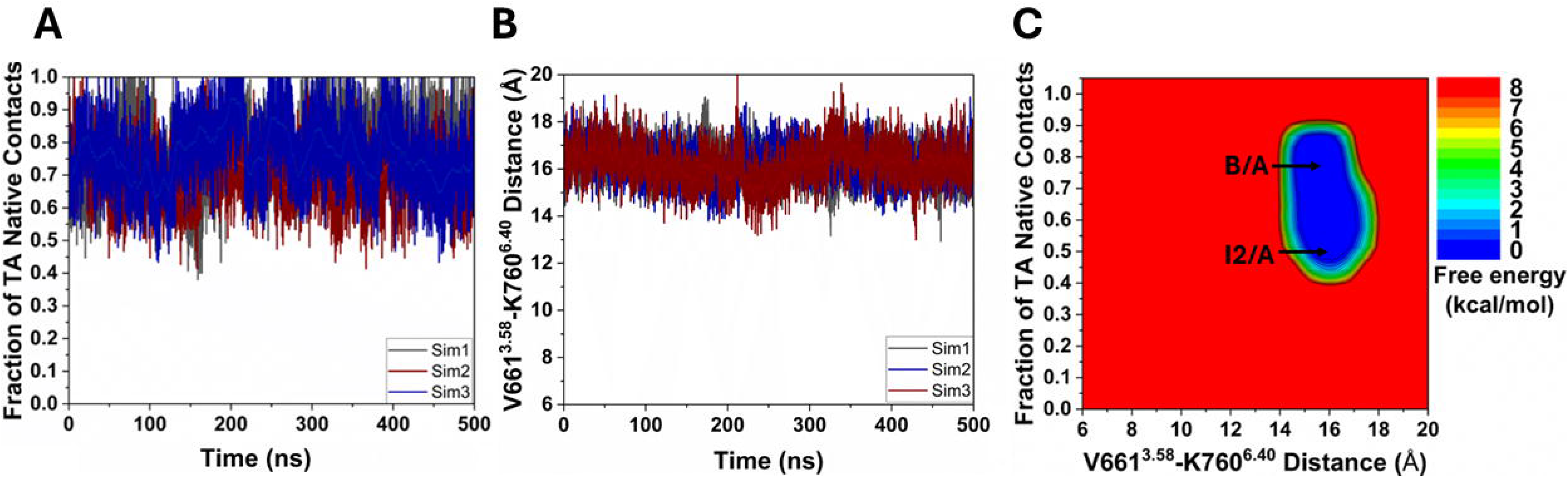

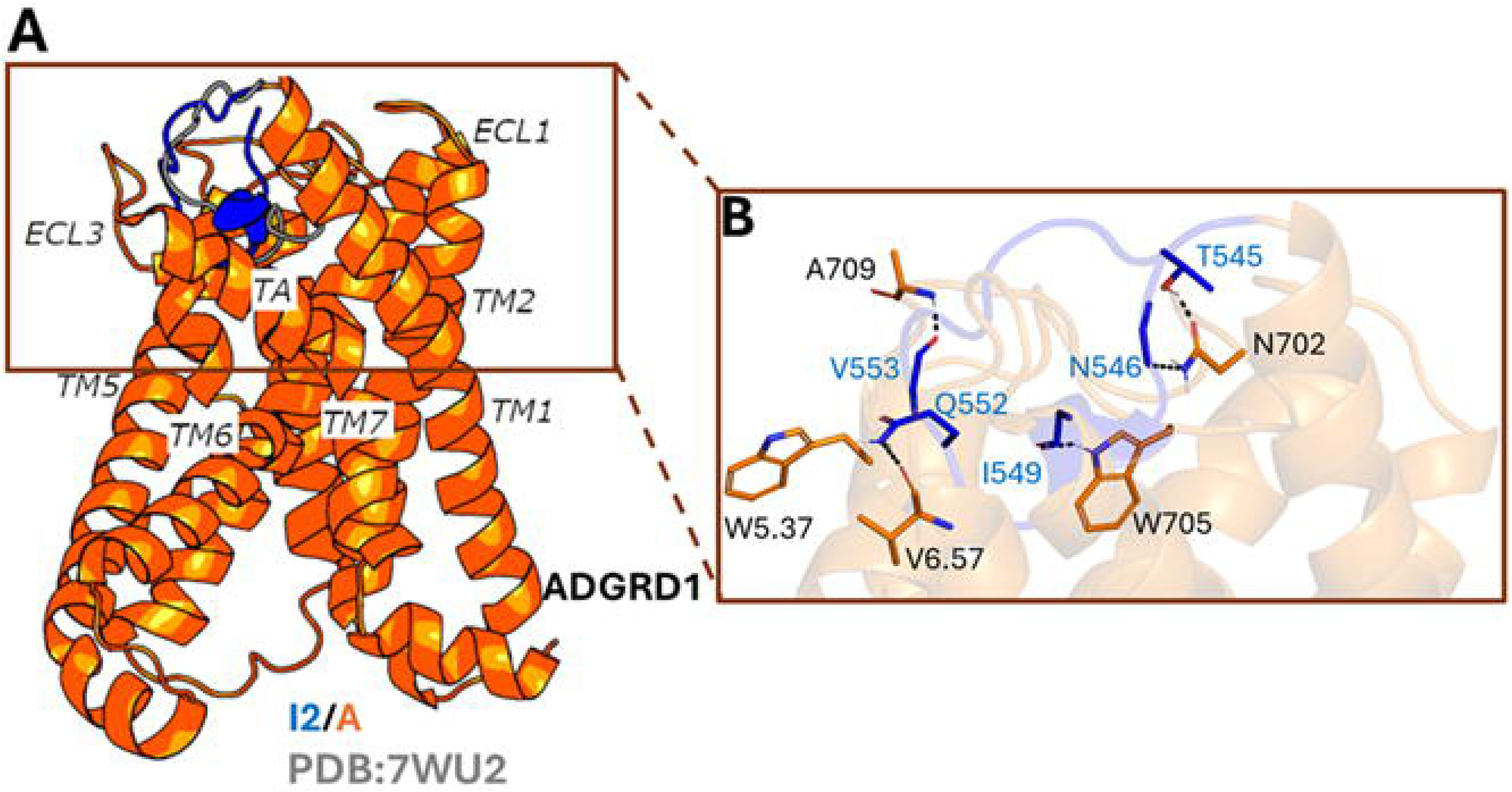

