## Supplementary material for "Mechanism of Tethered Agonist Binding to an Adhesion G-Protein-Coupled Receptor": The Supporting Information includes details about the Pep-GaMD method, Tables S1 to S4 and Figures S1 to S6

**Peptide-Gaussian accelerated Molecular Dynamics (Pep-GaMD)**

Based on Gaussian-accelerated Molecular Dynamics (GaMD)^1-4^, a new Pep-GaMD method^5^ is developed for improved sampling of peptide-protein interactions. With the new “Pep-GaMD” method^5^, we can simulate the repetitive binding and unbinding of peptides within microsecond simulations, which is challenging through cMD simulations^6, 7^.

Consider a system of peptide *L* binding to a protein target *P* in a biological environment *E*. The system consists of *N* atoms with their coordinates defined as $r\equiv\left\{ {\overset{⃑}{r}}_{1},\cdots,{\overset{⃑}{r}}_{N} \right\}$, and momenta defined as $p\equiv\left\{ {\overset{⃑}{p}}_{1},\cdots,{\overset{⃑}{p}}_{N} \right\}$.

The Hamiltonian of the system can be written as-

$H\left( r, p \right)=K\left( p \right)+V\left( r \right)$	(1)

where $K\left( p \right)$ and $V\left( r \right)$ denotes the kinetic and total potential energies of the system, respectively. The system’s potential energy term can further be divided into the following terms-

$V\left( r \right)=V_{P,b}\left( r_{P} \right)+V_{L,b}\left( r_{L} \right)+V_{E,b}\left( r_{E} \right)$

$$+ V_{PP,nb}\left( r_{P} \right)+V_{LL,nb}\left( r_{L} \right)+V_{EE,nb}\left( r_{E} \right)$$

$+V_{PL,nb}\left( r_{PL} \right)+V_{PE,nb}\left( r_{PE} \right)+V_{LE,nb}\left( r_{LE} \right)$	(2)

where $V_{P,b}$, $V_{L,b}$ and $V_{E,b}$ denotes the bonded potential energies of the protein *P*, peptide *L* and environment *E*, respectively. The self-non-bonded potential energies of protein *P*, peptide *L* and environment *E* are denoted as $V_{PP,nb}$, $V_{LL,nb}$ and $V_{EE,nb}$, respectively*.* The *P-L*, *P-E*, and *L-E's* related non-bonded interaction energies are denoted as $V_{PL,nb}$, $V_{PE,nb}$ and $V_{LE,nb}$, respectively.

According to classical molecular mechanics force fields^8^, the non-bonded potential energies are calculated as-

$V_{nb}=V_{elec}+V_{vdW}$	(3)

where $V_{elec}$ and $V_{vdW}$ represents the electrostatic and van der Waals potential energies of the system.

The binding process of peptides involves mainly their bonded and non-bonded interaction energies since peptides undergo large conformational changes during their binding to the target proteins. Therefore, the essential peptide potential energy is denoted as $V_{L}\left( r \right)= V_{LL,b}\left( r_{L} \right)+ V_{LL,nb}\left( r_{L} \right)+ V_{PL,nb}\left( r_{PL} \right)+ V_{LE,nb}\left( r_{LE} \right)$.

In Pep-GaMD^5^, we first add boost potential selectively to the essential peptide potential energy $V_{L}\left( r \right)$ based on the GaMD algorithm^1^-

${\Delta V}_{L}\left( r \right)=\left\{ \begin{aligned} \frac{1}{2}k_{L}\left( E_{L}-V_{L}\left( r \right) \right)^{2}, &V_{L}\left( r \right)<E_{L} \\ 0, &V_{L}\left( r \right)\geq E_{L} \end{aligned} \right.$	(4)

where E*_L_* denotes the threshold energy for applying the boost potential and *k_L_* represents the harmonic force constant. The Pep-GaMD^5^ simulation parameters are derived in a similar manner as shown in the previous GaMD methodology^1^.

When *E* is set to the lower bound with the system maximum potential energy (*E=V_max_*), the effective harmonic force constant$k_{0}$ can be written as-

$k_{0}=\min\left( 1.0, k_{0}^{'} \right)=min(1.0, \frac{\sigma_{0}}{\sigma_{V}}\frac{V_{max}-V_{min}}{V_{max}-V_{avg}})$,	(5)

where $V_{max}$, $V_{min}$, $V_{avg}$ and $\sigma_{V}$ are the maximum, minimum, average, and standard deviation of the boosted system potential energy, and $\sigma_{0}$ is the user-defined upper limit of the standard deviation of $\Delta V$ for proper reweighting. The harmonic force constant is denoted as $k=k_{0}\cdot\frac{1}{V_{max}-V_{min}}$ with ${0<k}_{0}\leq1$.

Alternatively, when the threshold energy *E* is set to its upper bound given as $E=V_{min}+\frac{1}{k}$, the effective harmonic force constant$k_{0}$ is set to-

$k_{0}=k_{0}^{"}\equiv(1-\frac{\sigma_{0}}{\sigma_{V}})\frac{V_{max}-V_{min}}{V_{avg}-V_{min}}$	(6)

if $k_{0}^{"}$ is found to be between 0 and 1. Otherwise,$k_{0}$ is calculated by using Eqn. (5).

In addition to selectively boosting the essential peptide potential energy, another boost potential is applied on the protein and solvent to increase the conformational sampling and facilitate peptide rebinding to the protein. The second boost potential is calculated using the total system potential energy other than the essential peptide potential energy as follows-

$\Delta V_{D}\left( r \right)=\left\{ \begin{aligned} \frac{1}{2}k_{D}\left( E_{D}-V_{D}\left( r \right) \right)^{2}, &V_{D}\left( r \right)<E_{D} \\ 0, &V_{D}\left( r \right)\geq E_{D} \end{aligned} \right.$	(7)

where *V_D_* is the total system potential energy (excluding the essential peptide potential energy), E_D_ corresponds to the threshold energy for applying the second boost potential and *k_D_* is the harmonic force constant, respectively. This leads to dual-boost Pep-GaMD algorithm^5^ where the total boost potential is given as $\Delta V\left( r \right)=\Delta V_{L}\left( r \right)+\Delta V_{D}\left( r \right)$.

**Energetic reweighting of Pep-GaMD for free energy calculations**

To perform energetic reweighting of Pep-GaMD^5^ simulations, the probability distribution defined along a selected reaction coordinate can be denoted as $p^{*}\left( A \right)$. Given $\Delta V\left( r \right)$ as the boost potential for each frame in Pep-GaMD simulations^5^, $p^{*}\left( A \right)$ can be reweighted to recover the canonical ensemble distribution, $p\left( A \right)$, as follows-

$p\left( A_{j} \right)=p^{*}\left( A_{j} \right)\frac{\left\langle e^{\beta\Delta V\left( \bar{r} \right)} \right\rangle_{j}}{\sum_{i=1}^{M} \left\langle{p^{*}\left( A_{i} \right)e}^{\beta\Delta V\left( \bar{r} \right)} \right\rangle_{i}}, j=1,\ldots, M$

(8)

where *M* denotes the number of bins, $\beta=k_{B}T$ and $\left\langle e^{\beta\Delta V\left( \bar{r} \right)} \right\rangle_{j}$ denotes the ensemble-averaged Boltzmann factor of $\Delta V\left( \bar{r} \right)$ for the simulation frames observed in the *j*^th^ bin. In order to reduce the energetic noise, the ensemble-averaged reweighting factor can be approximated using the cumulant expansion^9^-

$\left\langle e^{\beta\Delta V\left( \bar{r} \right)} \right\rangle=exp\left\{ \sum_{k=1}^{\infty} \frac{\beta^{k}}{k!}C_{k} \right\}$

(9)

where the first two cumulants are defined as-

$C_{1}= \left\langle\Delta V \right\rangle$

$$C_{2}= \left\langle\Delta V^{2} \right\rangle-\left\langle\Delta V \right\rangle^{2}=\sigma_{\Delta V}^{2}$$

(10)

The boost potential obtained from Pep-GaMD^5^ simulations usually shows near-Gaussian distribution. Therefore, cumulant expansion to the second order^9^ provides a more accurate reweighting. The reweighted free energy $F\left( A \right)={-k}_{B}T\ln p\left( A \right)$ is denoted as-

$F\left( A \right)=F^{*}\left( A \right)-\sum_{k=1}^{2} \frac{\beta^{k}}{k!}C_{k}+F_{c}$

(11)

where $F^{*}\left( A \right)={-k}_{B}T\ln p^{*}\left( A \right)$ denotes the modified free energy obtained from Pep-GaMD^5^ simulation and $F_{c}$ is a constant.

**Kinetic reweighting of Pep-GaMD simulations**

In Pep-GaMD^5^ simulations, reweighting of peptide dissociation and binding kinetics followed a similar protocol which is based on Kramer’s rate theory that has been implemented in kinetics reweighting of the GaMD methodology^1^. Given sufficient sampling of repetitive peptide dissociation and binding events in the simulations, we recorded the time periods for the peptide sampled in the bound (τ*_B_*) and unbound (τ*_U_*) states. The peptide binding and dissociation rate constants (*k*_off_ and *k*_on_) were calculated as-

$k_{off}=\frac{1}{\tau_{B}}$	(12)

$k_{on}=\frac{1}{\tau_{U} [L]}$	(13)

where [L] is the peptide concentration

Based on Kramers' rate theory, the rate of a chemical reaction in the large viscosity limit is calculated as^10^-

$k_{R}\cong\frac{w_{m}w_{b}}{2\pi\xi}e^{-{\Delta F}/{k_{B}T}}$	(14)

where $w_{m}$ and $w_{b}$ are frequencies of the approximated harmonic oscillators (also defined as curvatures of free energy surface^11^) near the energy minimum and barrier, respectively, $\xi$ is the frictional rate constant and $\Delta F$ is the free energy barrier of transition. The friction constant $\xi$ is related to the diffusion coefficient *D* with $\xi=k_{B}T/D$. The apparent diffusion coefficient *D* is calculated by dividing the kinetic rate obtained using the transition time series calculated directly from simulations by the probability density solution of the Smoluchowski equation^12^. In order to reweight peptide kinetics from the simulations using Kramer’s rate theory, the free energy barriers of peptide binding and dissociation are calculated from the original (reweighted, ***∆F***) and modified (no reweighting, ***∆F****) PMF profiles, similarly for curvatures of the reweighed (*w*) and modified ($w^{*}$, no reweighting) PMF profiles near the peptide bound (“B”) and unbound (“U”) low-energy wells and the energy barrier (“Br”), and the ratio of apparent diffusion coefficients from simulations without reweighting (modified, $D^{*}$) and with reweighting (*D*). The resulting numbers are then plugged into Eq. (14) to obtain accelerations of the peptide binding and dissociation rates during the Pep-GaMD^5^ simulations, which allows us to obtain the original kinetic rate constants.

**Table S1.** **Summary of the Pep-GaMD simulations performed on the human ADGRD1 in the presence of unbound tethered agonist (TA-unbound ADGRD1 system).**

| **System** | **System Size** | **ID** | **Simulation length** | **Boost Potential (kcal/mol)** |
| --- | --- | --- | --- | --- |
| **TA-ADGRD1** | 151,624 atoms | Sim1 | 1000ns | 18.62 ± 4.45 |
|  |  | Sim2 |  | 19.02 ± 4.46 |
|  |  | Sim3 |  | 19.21 ± 4.52 |
|  |  | Sim4 |  | 19.10 ± 4.44 |
|  |  | Sim5 |  | 19.33 ± 4.51 |

**Table S2.** **Summary of the Pep-GaMD simulations performed on the human ADGRD1-G_S_ in the presence of unbound tethered agonist (TA-unbound ADGRD1-G_S_ system).**

| **System** | **System Size** | **ID** | **Simulation length** | **Boost Potential (kcal/mol)** |
| --- | --- | --- | --- | --- |
| **TA-ADGRD1-Gs** | 249,391 atoms | Sim1 | 2500 ns | 40.64 ± 5.41 |
|  |  | Sim2 |  | 37.43 ± 5.35 |
|  |  | Sim3 |  | 39.85 ± 5.79 |
|  |  | Sim4 |  | 39.12 ± 5.42 |
|  |  | Sim5 |  | 37.65 ± 5.36 |

**Table S3.** **Summary of the Pep-GaMD simulations performed on the human ADGRD1 in the presence of bound tethered agonist (TA-bound ADGRD1 system).**

| **System** | **System Size** | **ID** | **Simulation length** | **Boost Potential (kcal/mol)** |
| --- | --- | --- | --- | --- |
| **TA-ADGRD1** | 109,011 atoms | Sim1 | 500 ns | 12.74 ± 3.94 |
|  |  | Sim2 |  | 12.24 ± 3.90 |
|  |  | Sim3 |  | 12.76 ± 3.95 |

**Table S4.** **Summary of the Pep-GaMD simulations performed on the human ADGRD1-G_S_ in the presence of bound tethered agonist (TA-bound ADGRD1-G_S_ system).**

| **System** | **System Size** | **ID** | **Simulation length** | **Boost Potential (kcal/mol)** |
| --- | --- | --- | --- | --- |
| **TA-ADGRD1-Gs** | 159,372 atoms | Sim1 | 500 ns | 13.28 ± 4.04 |
|  |  | Sim2 |  | 12.38 ± 3.87 |
|  |  | Sim3 |  | 12.52 ± 3.90 |

**Figure S1.** Simulation starting structure of the **(A)** ADGRD1-TA (unbound) complex with G_S_ protein added. **(B)** ADGRD1-TA (bound) complex with G_S_ protein added. **(C)** ADGRD1-TA (unbound) complex with G_S_ protein removed. **(D)** ADGRD1-TA (bound) complex with G_S_ protein removed.

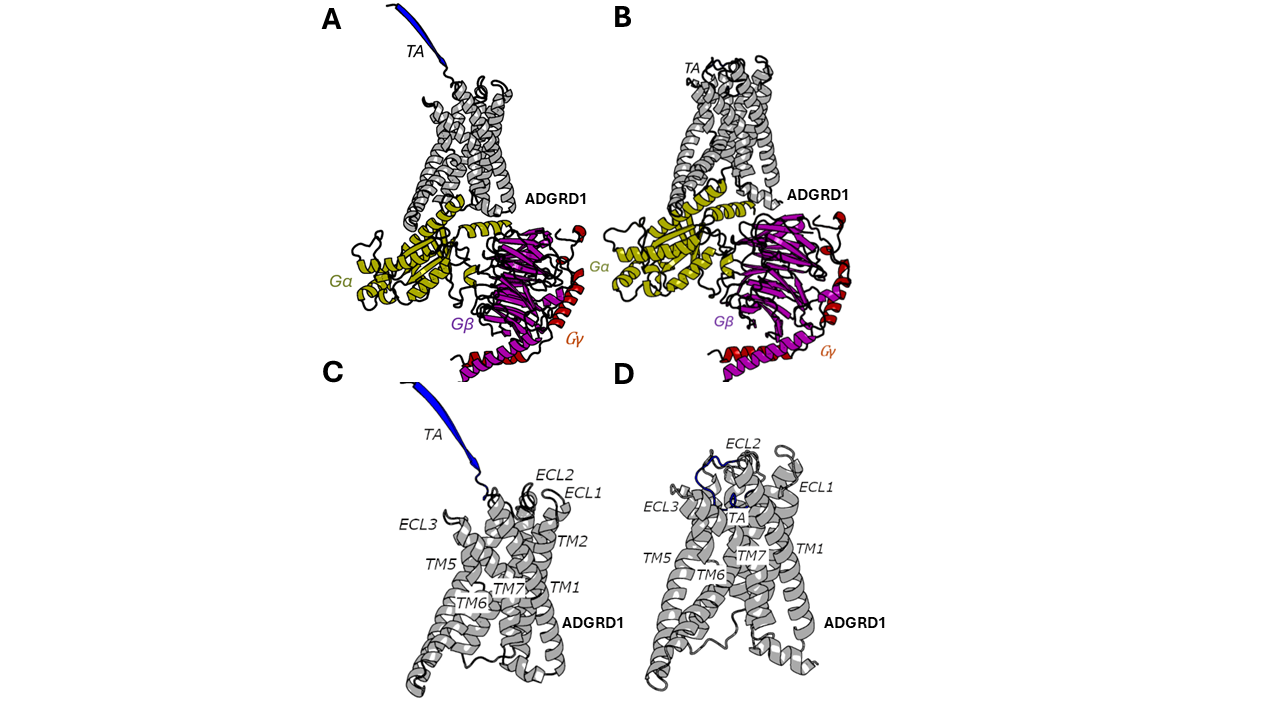

**Figure S2.** Computational model of the **(A)** ADGRD1-TA (unbound) complex with G_S_ protein added and **(B)** ADGRD1-TA (unbound) complex with G_S_ protein removed. The systems were embedded in membrane lipids and solvated in aqueous medium. The phosphatidylcholine (POPC) membrane lipids were rendered as cyan sticks, the ADGRD1 receptor as grey cartoons, and the sodium and chlorine ions as yellow and green spheres.

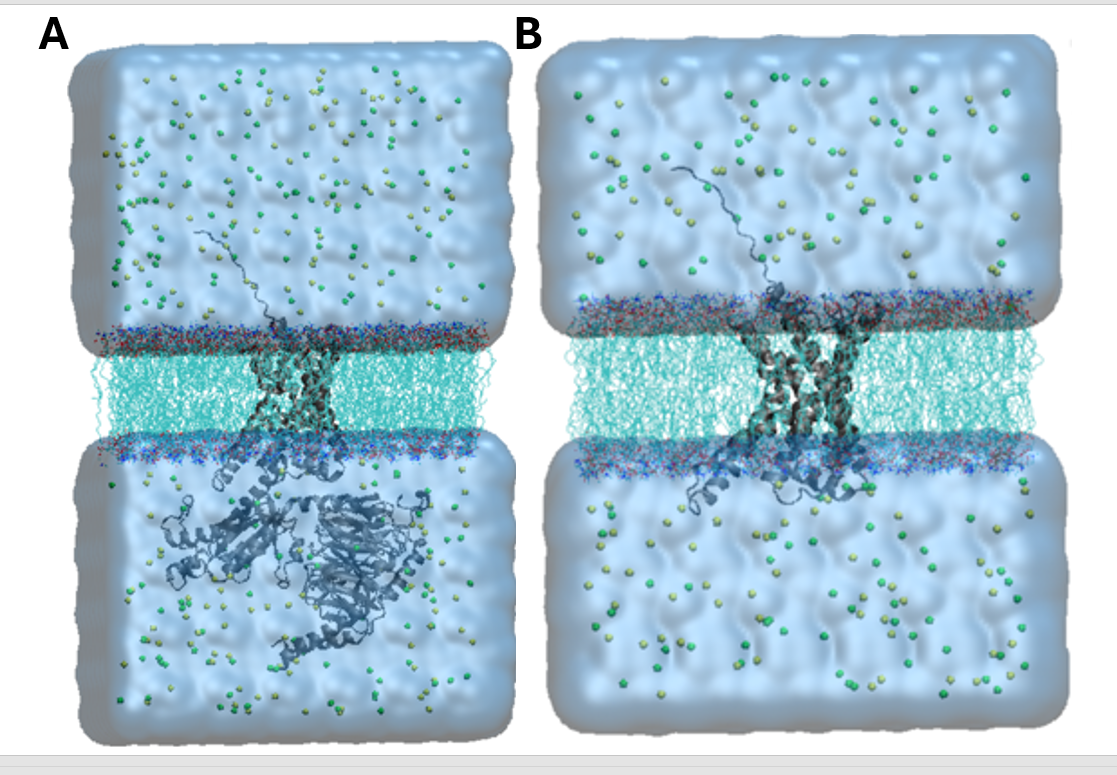

**Figure S3. Binding of tethered agonist (TA) in the G_S_ protein-coupled ADGRD1 receptor was observed in Pep-GaMD simulations:** **(A)** Time course of the distance between the Cα atoms of V661^3.58^ and K760^6.40^ residues calculated from five 2500 ns Pep-GaMD simulations. **(B)** 2D potential of mean force (PMF) free energy profile regarding the fraction of native contacts between the TA and receptor and V661^3.58^-K760^6.40^ distance calculated by combining five 2500 ns Pep-GaMD simulations. The low-energy states are labeled as “Unbound/Active” (U/A) and “Bound/Active” (B/A).

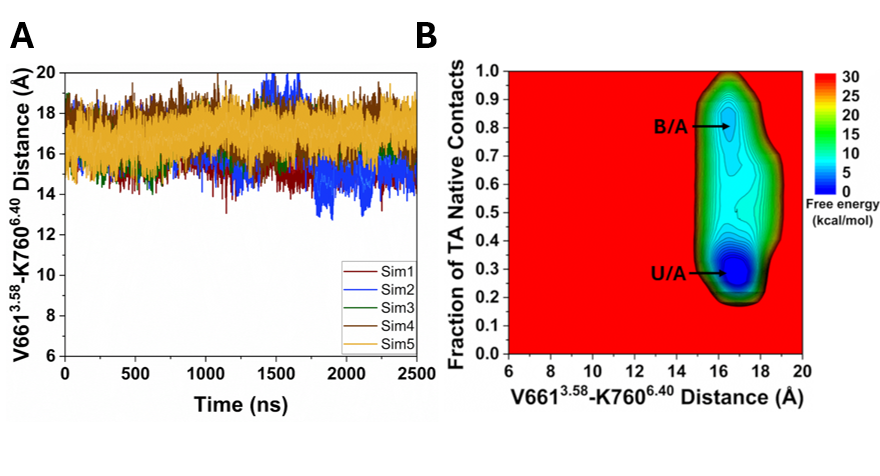

**Figure S4.** **Dynamic motions of bound tethered agonist (TA) in the ADGRD1-G_S_ protein complex were observed in Pep-GaMD simulations: (A)** Time course of the distance between the Cα atoms of V661^3.58^ and K760^6.40^ residues calculated from three 500 ns Pep-GaMD simulations. **(B)** 2D potential of mean force (PMF) free energy profile regarding the fraction of native contacts between the TA and receptor and V661^3.58^-K760^6.40^ distance calculated by combining three 500 ns Pep-GaMD simulations. The low-energy states are labeled as “Intermediate2/Active” (I2/A) and “Bound/Active” (B/A).

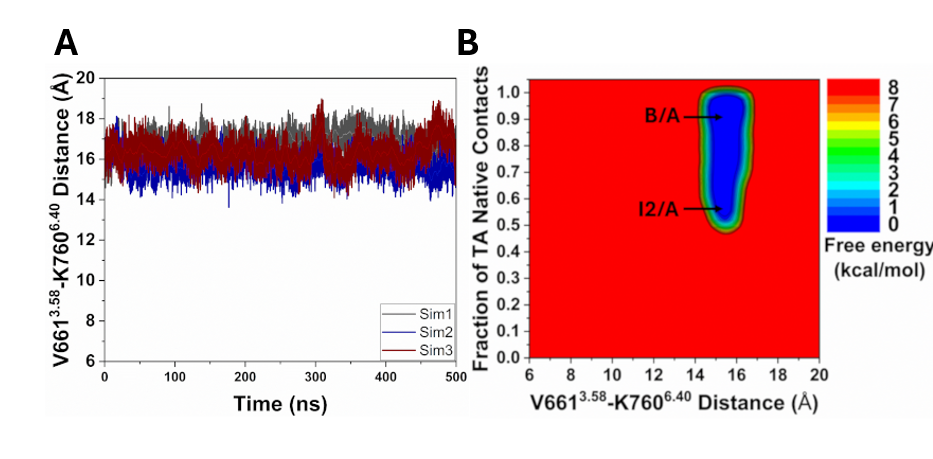

**Figure S5. Dynamic motions of bound tethered agonist (TA) in the ADGRD1 receptor were observed in Pep-GaMD simulations: (A)** Time course of the fraction of native contacts between the TA and receptor calculated from three 500 ns Pep-GaMD simulations. **(B)** Time course of the distance between the Cα atoms of V661^3.58^ and K760^6.40^ residues calculated from three 500 ns Pep-GaMD simulations. **(C)** 2D potential of mean force (PMF) free energy profile regarding the fraction of native contacts between the TA and receptor and V661^3.58^-K760^6.40^ distance calculated by combining three 500 ns Pep-GaMD simulations. The low-energy states are labeled as “Intermediate 2/Active” (I2/A) and “Bound/Active” (B/A).

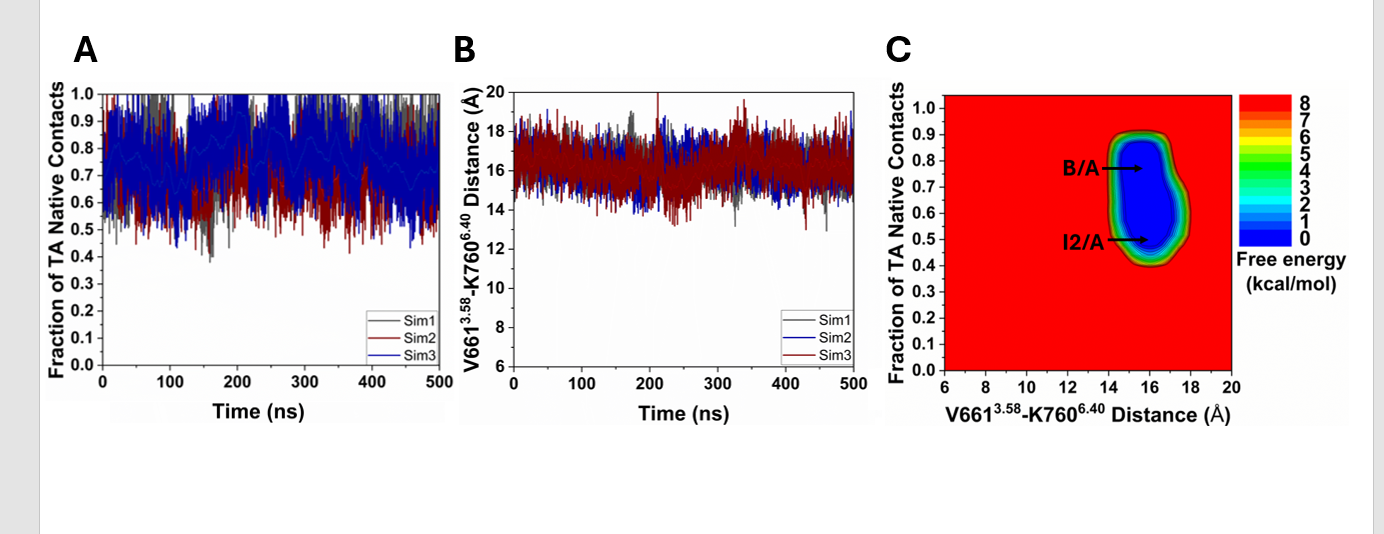

**Figure S6. (A)** The low-energy conformation of the TA-bound ADGRD1 complex in the “Intermediate 2/Active” (I2/A) state compared with the cryo-EM structure (grey, PDB: 7WU2). **(B)** Critical interactions between TA (blue) and ADGRD1 (orange) observed in the I2/A state. The tethered agonist formed polar and hydrophobic interactions with receptor residues N702^ECL2^, W705^ECL2^, A709^ECL2^, W^5.37^ and V^6.57^.

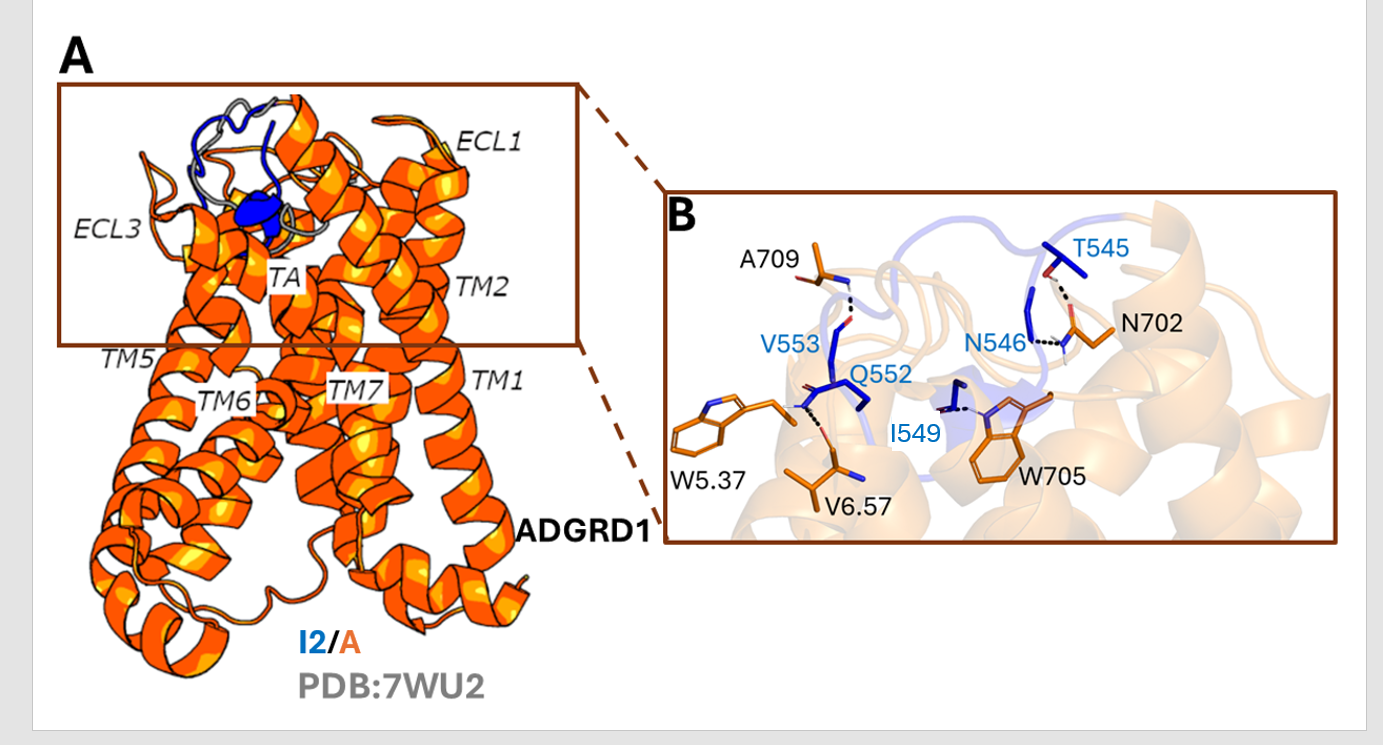
